# Quantifying uncertainty of taxonomic placement in DNA barcoding and metabarcoding

**DOI:** 10.1101/070573

**Authors:** Panu Somervuo, Douglas Yu, Charles Xu, Yinqiu Ji, Jenni Hultman, Helena Wirta, Otso Ovaskainen

## Abstract

1. A crucial step in the use of DNA markers for biodiversity surveys is the assignment of Linnaean taxonomies (species, genus, etc.) to sequence reads. This allows the use of all the information known based on the taxonomic names. Taxonomic placement of DNA barcoding sequences is inherently probabilistic because DNA sequences contain errors, because there is natural variation among sequences within a species, and because reference databases are incomplete and can have false annotations. However, most existing bioinformatics methods for taxonomic placement either exclude uncertainty, or quantify it using metrics other than probability.
2. In this paper we evaluate the performance of a recently proposed probabilistic taxonomic placement method PROTAX by applying it to both annotated reference sequence data as well as unknown environmental data. Our four case studies include contrasting taxonomic groups (fungi, bacteria, mammals, and insects), variation in the length and quality of the barcoding sequences (from individually Sanger-sequenced sequences to short Illumina reads), variation in the structures and sizes of the taxonomies (from 800 to 130 000 species), and variation in the completeness of the reference databases (representing 15% to 100% of the species).
3. Our results demonstrate that PROTAX yields essentially unbiased assessment of probabilities of taxonomic placement, and thus that its quantification of species identification uncertainty is reliable. As expected, the accuracy of taxonomic placement increases with increasing coverage of taxonomic and reference sequence databases, and with increasing ratio of genetic variation among taxonomic levels over within taxonomic levels.
4. Our results show that reliable species-level identification from environmental samples is still challenging, and thus neglecting identification uncertainty can lead to spurious inference. A key aim for future research is the completion and pruning of taxonomic and reference sequence databases, and making these two types of data compatible.

## Introduction

In this paper, we use the term ‘DNA barcoding’ to refer to molecular species identification with the help of ‘barcoding’ genes, which are short sequences of DNA that vary greatly between species but little within species (Hebert et al. 2003). DNA barcoding has revolutionized biological studies by increasing the speed and reliability of assigning Linnaean taxonomies to biological specimens (Ratnasingham and Hebert 2007). When combined with high-throughput sequencing, barcoding can be applied to bulk samples or environmental DNA, which approach we call here ‘DNA metabarcoding’ (Taberlet et al. 2012; Yu et al. 2012).

In the metabarcoding pipeline, DNA is extracted from a bulk sample containing potentially multiple species, a taxonomically informative gene is PCR-amplified, and the resulting PCR-products are sequenced. The raw sequence output is processed through a bioinformatics pipeline that includes denoising and removal of low quality and chimeric sequences, assignment of sequences to their samples, and grouping similar sequences into ‘operational taxonomic units’ (OTUs). OTUs are meant to represent distinct biological taxa, usually distinct species. The term OTU indicates that the clusters are not necessarily biological species but that they can be considered as species hypotheses. This is because OTUs are typically defined phenetically using a sequence-similarity threshold. Finally, in a crucial step, the researcher wishes to know the species identities behind the OTUs, i.e. to place them into a Linnaean taxonomy.

Taxonomic placement of OTUs to high-level ranks (phylum, class, order) is relatively straightforward (e.g. Yu et al. 2012), whereas placement to lower ranks (family, genus, species) has remained more difficult. This is partly because of the limited information contained in the short sequences generated by high-throughput sequencing platforms, and partly because of the incomplete nature of reference databases, with missing taxa and limited within-taxon sampling (Lou and Golding 2012). Furthermore, widely applied methods for low-level taxonomic placement lack a proper assessment of identification reliability. For example, a user of the Barcode of Life Database System (www.boldsystems.org, accessed 5 Aug 2016) encounters the warning “this search only returns a list of the nearest matches and does not provide a probability of placement to a taxon”. As we discuss in more detail below, the ability to conduct reliable low-level taxonomic placement would make major contributions to species-level analyses, community-level analyses, as well as metabarcoding methodology itself.

The value of assigning species names to barcoding sequences is that it allows one to link the samples to the rest of our vast biological knowledge (Janzen et al. 2005). For instance, if mammalian DNA isolated from a mosquito blood meal can be reliably assigned to red fox (*Vulpes vulpes*), it enables one to combine the sample with many other kinds of information. These may include information on the red fox's behaviour, population growth rate, age structure, geographic distribution, habitat requirements, and trophic position, such as its top-down control of rodent vectors of Lyme disease (Levi et al. 2012). More generally, accurate low-level taxonomic placement of metabarcoding sequences improves many kinds of assessments of the structure and function of communities, and how these change over space, time, or environmental gradients. For example, species-level identifications of gut contents or faeces allows the construction of high-resolution food webs (e.g. Wood et al. 2015). As another example, environmental change can be inferred through species-level identification of ancient DNA, derived e.g. from lake sediments (Pansu et al. 2015). As a further example, in food and medicine, DNA barcoding can be used to improve food safety and wildlife forensics (Staats et al. 2016), e.g. through the detection of falsely labelled products (Wong and Hanner 2008) and forbidden ingredients (Coghlan et al. 2012). As a final example, metabarcoding can be used to monitor nature reserves and to detect endangered species, e.g. rare rainforest mammals from the residual blood meals of leeches (Schnell et al. 2012).

The ability to conduct accurate low-level taxonomic placement would also contribute to the metabarcoding methodology itself. Although OTUs are meant to represent single species, biological species can unintentionally be split or merged during OTU clustering. Accurate species-level taxonomic placement enables one to merge multiple OTUs that receive identical taxonomic placements. Conversely, cases in which a single OTU is assignable with equal confidence to multiple species can be used to identify taxonomic groups that would benefit the most from better reference databases or where taxonomic revision may be needed. In addition, accurate low-level taxonomic placement makes it easier to detect and remove contaminant OTUs.

As demonstrated by the above examples, for many kinds of purposes it is of critical importance to know when we can and when we cannot reliably identify an OTU down to family, genus, or species. Many kinds of bioinformatics programs are currently available for taxonomic placement. These can be classified into three general categories: similarity-based, similarity/phylogeny-based, and phylogenetic-placement-based. The most common are those that compare the similarities between the environmental sequences and the sequences of the reference database: BLAST (Altschul et al. 1997), MEGAN (Huson et al. 2007), BOLD (Ratnasingham and Hebert 2007), UTAX (Edgar 2013), NBC (Wang et al. 2007), and the Geneious Sequence Classifier (Kearse et al. 2012). Similarity-based methods find the most phenetically similar reference sequence, and they do not thus define taxonomic clades based on fundamental principles from systematic biology related to synapomorphies (shared, derived characters). The second category of similarity/phylogeny-based methods is represented by the Statistical Assignment Program (SAP), which first uses BLAST to create a group of sequence homologues for an OTU (Munch et al. 2008). Multiple phylogenetic trees are then generated for the OTU and its homologues, and taxonomic placement is guided by the summarised position of the OTU within the trees. The third category of phylogenetic-placement-based methods includes *pplacer* (Matsen et al. 2010) and the Evolutionary Placement Algorithm (EPA) (Berger et al. 2011). These methods first construct a single maximum-likelihood phylogeny from all available reference sequences, after which they place the OTUs within the phylogenetic tree.

A major challenge affecting all taxonomic placement methods is that reference databases are incomplete, and that they may contain mislabelled reference sequences. This is especially problematic when trying to identify a sequence within a large taxonomic clade in regions of high biodiversity where many organisms have yet to be sequenced. Ideally, uncertainty due to incomplete or mislabelled reference sequences should result in taxonomic placement to higher taxonomic ranks, not to the most similar reference sequence that happens to be available. Thus far, only heuristic solutions to this problem have been proposed. For example, in MEGAN, a lowest-common-ancestor (LCA) assignment algorithm uses several best BLAST hits to determine the taxonomic level into which the assignment is given, but incomplete reference databases may still lead to false annotations.

In our previous work, we developed the bioinformatics pipeline PROTAX (PRObabilistic TAXonomic placement, Somervuo et al. 2016) which accounts explicitly for incompleteness of taxonomic and reference databases. This is achieved by placing environmental sequences into a Linnaean taxonomy that is typically only partly populated by reference sequences. The taxonomic placements generated by PROTAX include known taxonomic units (species present in the Linnaean taxonomy) for which reference sequences are available, known taxonomic units for which reference sequences are not available, and unknown taxonomic units, such as species or genera that are missing from the Linnaean taxonomy. A key feature of PROTAX is that it is probabilistic, i.e. it decomposes the probability of one among all possible assignment outcomes. In the ideal case, one of the outcomes obtains a high probability whereas the other taxonomic placements obtain probabilities close to zero. In ambiguous cases, several outcomes obtain a non-negligible classification probability, and thus reliable taxonomic placement can be achieved only at a higher taxonomic rank. PROTAX is based on a statistically rigorous model, making its classification probabilities unbiased, as shown in Somervuo et al. (2016) for simulated data and a small-scale empirical case study. In other words, if PROTAX assigns an 80% probability for placement to a given taxonomic unit for 100 sequences, the classification will be on average correct for 80 of those sequences, whereas it will not be correct for 20 of the sequences.

This paper has two aims. The first aim is to evaluate the potential of DNA (meta)barcoding for obtaining species-level identifications, given the current state of taxonomic databases, sequence reference databases, and sequencing technologies. The second aim is to evaluate the performance of PROTAX as a general tool for taxonomic placement. To address both aims, we apply PROTAX to four contrasting case studies, which differ greatly in their taxonomic scope (fungi, insects, mammals, and bacteria), the number of species involved, the coverage and quality of the reference databases, and the sequencing technology applied to environmental data. For each case study, we conduct two kinds of analyses. First, we examine how well PROTAX is able to classify validation sequences sampled from the reference database. Second, we apply PROTAX to environmental sequence data to examine the level of species identification resolution that can be expected to be achieved by different kinds of empirical studies.

## Materials and methods

We consider four case studies, for each of which we use three kinds of data: a taxonomy database, a reference sequence database, and environmental sequences originating from an empirical study (Table 1). The case studies vary greatly in many aspects: their taxonomic scopes (mammals, fungi, insects and bacteria), the sizes and coverages of the taxonomies and the reference databases, the barcoding gene used, and the sequencing technology applied. These influence e.g. the level of overlap among genetic variation between consecutive taxonomic levels (Fig. 1), with obvious implications to the possibility of species-level taxonomic placement. As the four case studies vary simultaneously in many aspects, their comparison does not enable asking e.g. whether it is generally easier to identify insects or fungi. Instead, they are selected to be diverse in order to illustrate the many kinds of issues that influence the accuracy of taxonomic placement.

**Figure 1.**
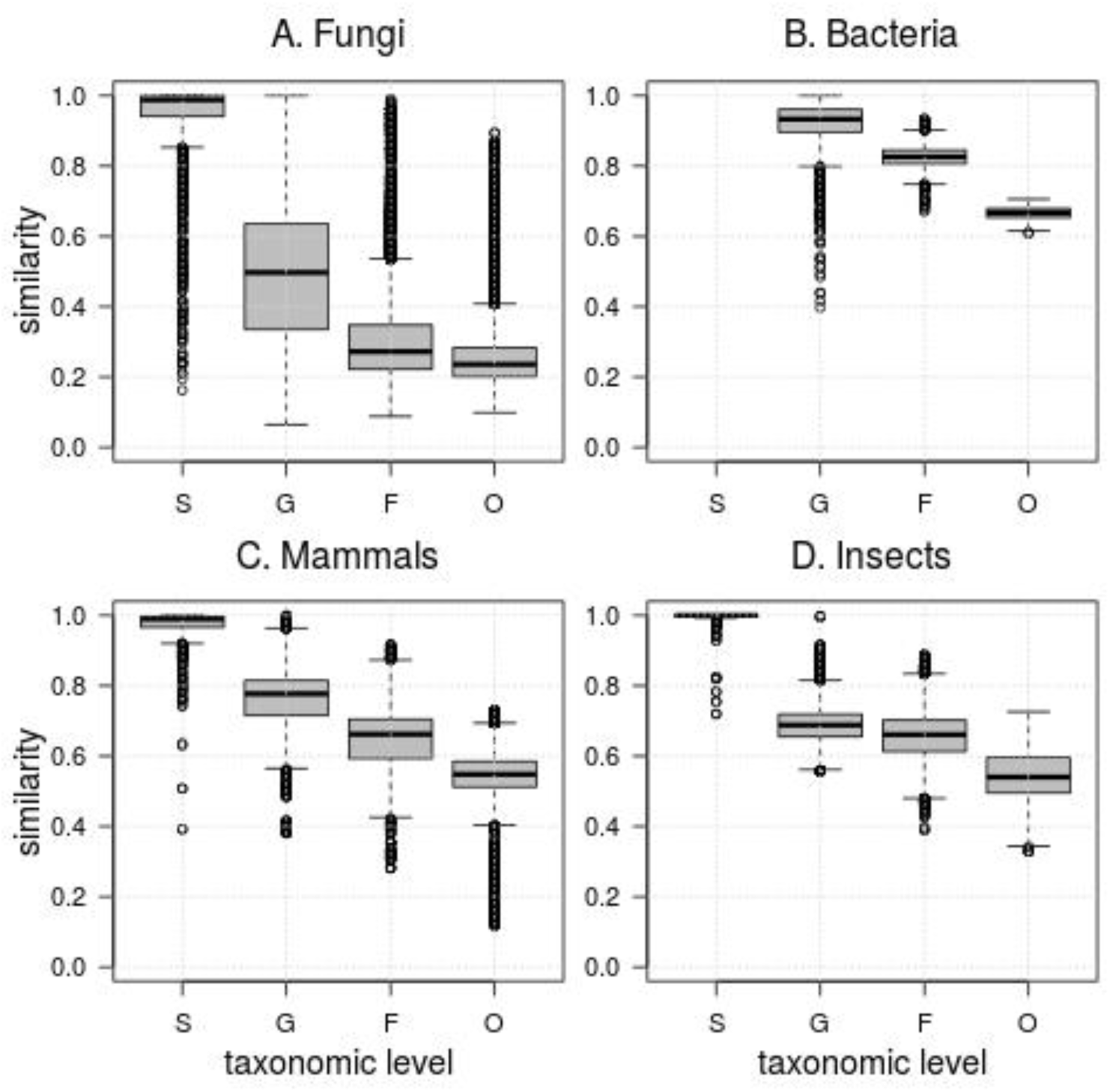
The distribution of pairwise LAST-similarities between reference sequences within each taxonomical levels of species (S), genus (G), family (F) and order (O). The distribution of similarities in a given taxonomical level originates from 1000 randomly selected sequence pairs. At the species level, each sequence pair represents two different individuals of the same species. At the genus, family, and order levels, each sequence pair represents, respectively, two different species, genera, or families that belong to the same genus, family, or order.

**Table 1.**
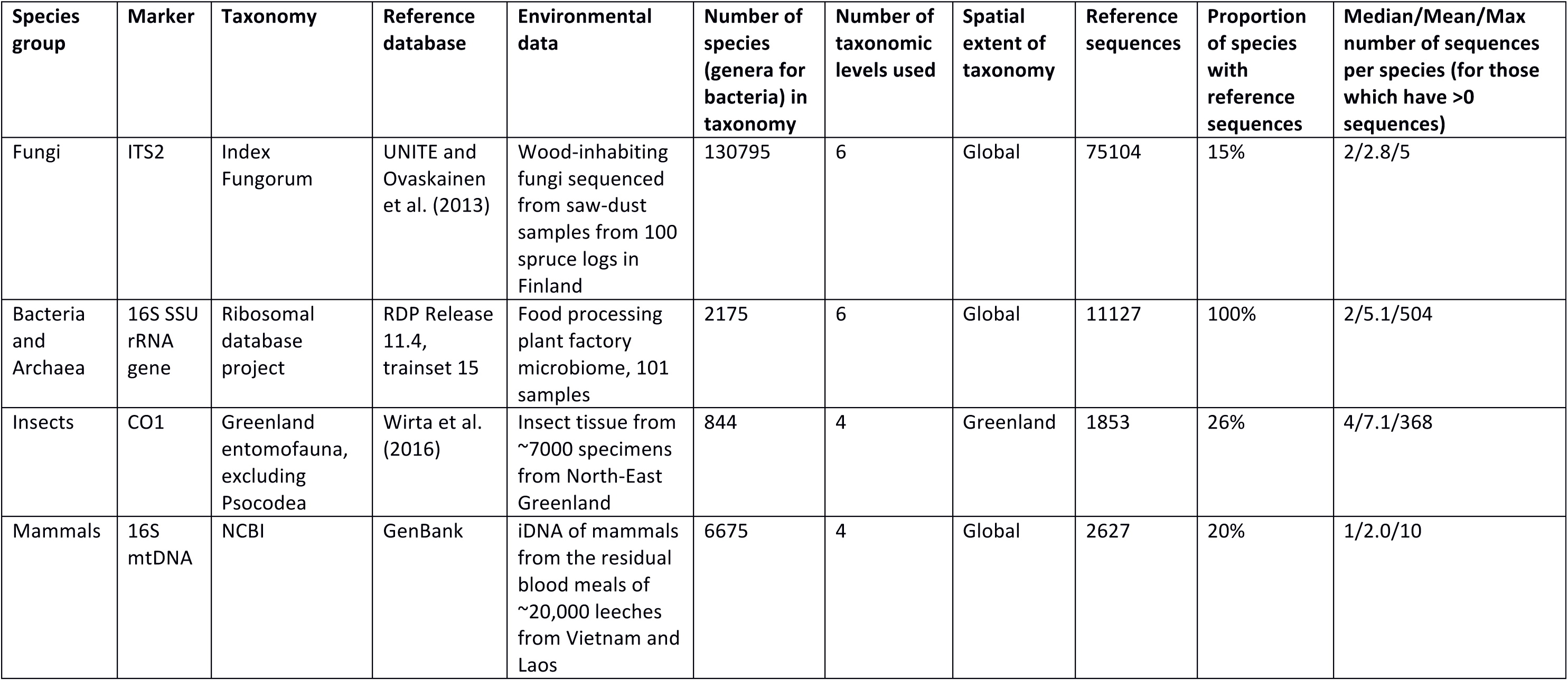
Case studies used to evaluate the performance of PROTAX in probabilistic taxonomic placement of environmental sequence data.

| Species group | Marker | Taxonomy | Reference database | Environmental data | Number of species (genera for bacteria) in taxonomy | Number of taxonomic levels used | Spatial extent of taxonomy | Reference sequences | Proportion of species with reference sequences | Median/Mean/Max number of sequences per species (for those which have >0 sequences) |
| --- | --- | --- | --- | --- | --- | --- | --- | --- | --- | --- |
| Fungi | ITS2 | Index Fungorum | UNITE and Ovaskainen et al. (2013) | Wood-inhabiting fungi sequenced from saw-dust samples from 100 spruce logs in Finland | 130795 | 6 | Global | 75104 | 15% | 2/2.8/5 |
| Bacteria and Archaea | 16S SSU rRNA gene | Ribosomal database project | RDP Release 11.4, trainset 15 | Food processing plant factory microbiome, 101 samples | 2175 | 6 | Global | 11127 | 100% | 2/5.1/504 |
| Insects | CO1 | Greenland entomofauna, excluding Psocodea | Wirta et al. (2016) | Insect tissue from ~7000 specimens from North-East Greenland | 844 | 4 | Greenland | 1853 | 26% | 4/7.1/368 |
| Mammals | 16S mtDNA | NCBI | GenBank | iDNA of mammals from the residual blood meals of ~20,000 leeches from Vietnam and Laos | 6675 | 4 | Global | 2627 | 20% | 1/2.0/10 |

For each case study, we first utilized the taxonomy and reference sequence databases to parameterize the PROTAX statistical model. To do so, we followed Somervuo et al. (2016), except for small modifications that we describe below. We then used the parameterized model to classify a set of well-identified reference sequences, with the aim of evaluating the classification accuracy of PROTAX at different taxonomic levels, and to assess if the classification probabilities are unbiased. Finally, we clustered the environmental data to OTUs, roughly at the species level, picked the most common sequence to represent each cluster, and used the parameterized PROTAX model for probabilistic taxonomic placement of these OTUs. The aim here was to assess how large a fraction of environmental data can be reliably classified to each taxonomic level, and to examine which fraction of environmental sequence data represents the two unknown categories included in PROTAX: species that are present in the taxonomy but for which reference sequences are available, and species that are missing from the taxonomy.

We first describe the three data types (taxonomy database, reference database, and environmental data) that we acquired for each case study, as well as make some remarks about the particularities of each case study. We then explain how PROTAX was fit to these data and how we assessed PROTAX's performance in probabilistic taxonomic placement.

### Identifying mammals from leech blood meals

#### Taxonomy database

We used the NCBI taxonomy (NCBI Resource Coordinators 2016) of all clades within Mammalia. This database has high coverage as it includes all 6674 species for which molecular data are available. The taxonomy is classified to the four levels of order, family, genus and species. For some species, classifications to intermediate levels or species-level were missing.

#### Reference sequence database

We used all available mammalian mitochondrial 16S rRNA gene sequences (mt 16S rRNA) downloaded from GenBank (Clark et al. 2016). We removed ambiguous bases and kept only sequences of length 300-1600 bp. We included at most 10 sequences of per species, resulting in a database of 2627 sequences representing 1315 different species.

#### Environmental sequences

We used mammalian mt 16S rRNA gene sequences (see Schnell et al. 2012 for further details on primer) derived from residual blood meals of ~20,000 haematophagous leeches collected in the central Annamite mountains of Vietnam and Laos (Yu et al., unpublished data). DNA extraction was conducted using the Qiagen QIAquick PCR purification kit, and sequencing by Illumina HiSeq 2000. Raw reads were denoised with *bfc* (Li 2015), chimeras removed by UCHIME (Edgar et al. 2011), and the sequences assigned to samples using the QIIME (Caporaso et al. 2010) script split_libraries.py. The reads were clustered into OTUs at 98% similarity using CROP (Hao et al. 2011). OTU*s* that were not identified as vertebrate mt 16S rRNA based on BLAST against GenBank were removed.

#### Remarks

As we use here individual, very short single-read (typically 100 bp) sequences provided by Illumina HiSeq, we aim to demonstrate how PROTAX performs in the case of high identification uncertainty, rather than attempting to identify the specimens as well as would be possible e.g. by including an assembly step. To illustrate the effect of sequence length, we parameterized the model both for full length and short length sequences.

### Identifying insects from individually sequenced specimens

#### Taxonomy database

We compiled a list of all species of the class Insecta (excluding Psocodea) recorded in Greenland, based on Böcher et al. 2015, with additions from Wirta et al. (2016). The environmental sequence data (see below) comes from the same study region as that of Wirta et al. (2016), and thus the taxonomic database is expected to cover the species of the region relatively well. The 1332 taxa were classified to the four levels of order, family, genus and species. Most of the taxa were defined to species, but a fraction as a sole representative of a genus.

#### Reference sequence database

We used barcode sequences of specimens collected from Zackenberg, Greenland. The reference database included the standard cytochrome *c* oxidase 1 (CO1) barcode sequence for 241 morphologically identified insect species (deposited in BOLD under dataset dx.doi.org/10.5883/DS-ZACKANIM).

#### Environmental sequences

We used 7939 CO1 sequences from insect tissue caught on sticky traps mimicking a flower in northeast Greenland. Each sequence (deposited in BOLD with the code ZACKD) represents a separate specimen (Tiusanen et al., unpubl. data) that was Sanger sequenced in one direction.

#### Remarks

As here both the taxonomy database as well as the reference database are specifically tailored to the environmental data, and as here the environmental sequence data consist of high quality sequences, this case study is aimed to illustrate a best case scenario.

### Identifying wood-inhabiting fungi from saw dust samples

#### Taxonomy database

We used the Index Fungorum database (www.indexfungorum.org), classified into the six levels of phylum, class, order, family, genus, and species. We reduced the amount of redundancy in the taxonomy by removing likely synonyms, such as old names of species that had been renamed. The resulting taxonomy consists of 130 795 species.

#### Reference sequence database

To construct the reference database of 75 104 sequences, we used the UNITE+INSD sequence database (https://unite.ut.ee/) consisting of fungal ITS region, complemented with the database of Ovaskainen et al. (2013). In order to increase the coverage of the reference sequences for poorly studied species groups, we also included those species hypothesis (SH) from UNITE that were more than 97% divergent from the other reference sequences. We extracted the ITS2 region of the reference sequences using ITSx software (Bengtsson-Palme et al. 2013). The majority (73%) of the reference sequences were annotated to the species level, but many only to the genus (11%) level or family or higher levels (16%). We included at most five sequences per species.

#### Environmental sequences

We used fungal ITS2 sequences originating from the study of Ovaskainen et al. (2013). The saw dust samples originate from 100 spruce logs sampled in autumn 2008 in a natural forest in southern Finland. DNA extraction was conducted using the Power Soil DNA isolation kit (MoBio Laboratories, Inc., Carlsbad, CA, USA), and sequencing was done on a Genome Sequencer FLX (454 Life Sciences, Roche, Branford, CT, USA). We removed all sequences that were shorter than 150 bp, resulting in 259 327 sequences. We used cutadapt (Martin 2011) to detect the presence of ITS4 primer in order to be sure that the sequence represented ITS2 region. To cope with homopolymer errors, all consecutive repetitions of the same nucleotide were removed as in Ovaskainen et al. (2010, 2013), both for reference and environmental sequences. Environmental sequences were clustered using UCLUST (Edgar 2010) with 99% identity threshold.

#### Remarks

This case study is aimed to illustrate how PROTAX copes with a very large taxonomy that is only poorly covered by reference sequences. We further use the fungal case study to examine how additional information can be incorporated into the PROTAX model: in addition to the baseline model, we constructed an alternative model, where we gave more weight to species that are expected to be found from the geographic area where the sampling was conducted (for more details, see below).

### Identifying bacteria from a food production pipeline

#### Taxonomy database

The taxonomy used for bacteria is different from other taxonomies in the sense that it is not an independent Linnaean taxonomy but it was generated from the Ribosomal Database Project (RDP) reference sequences (Wang et al. 2007) and therefore fully coincides with the reference sequence database (see below). The RDP reference taxonomy contains both bacteria and archaea, and it is well curated down to the genus level. Here we included the six levels of domain, phylum, class, order, family and genus. The taxonomy consists of 60 phyla classified to 2175 genera.

#### Reference sequence database

For the reference sequence database, we used the RDPClassifier training sequences, labeled to the genus level (trainset15_092015.fa from RDPClassifier_16S_trainsetNo15_rawtrainingdata.zip, available at rdp.cme.msu.edu/misc/resources.jsp).

#### Environmental sequences

We used bacterial 16S rRNA gene sequences from the study of Hultman et al. (2015), who aimed to understand the effect of food-preparation-surface microbiomes on the end product. The samples originate from surfaces of a food processing facility, from raw food material, and from cooked food products. As detailed in Hultman et al. (2015), total DNA was extracted from the samples using a bead beating method. The V1 to V3 region was PCR-amplified and sequenced with 454 GS FLX. The reads were quality filtered, chimeras were removed, and reads were assigned to OTUs with QIIME (Caporaso et al. 2010) using 97% similarity. Homopolymers were treated as in the fungal data. The raw sequence reads can be downloaded from Sequence Read Archive (SRA) of the NCBI under BioProject number PRJNA293141.

#### Remarks

As noted above, the bacterial case study differs fundamentally from the other case studies as the taxonomy database is not independent of the reference sequence database. Compared especially to mammals and Greenland insects, the taxonomy is likely to be incomplete. Thus with this case study we were interested in examining whether the environmental sample includes a high fraction of material that PROTAX would classify to belong to missing branches.

### Fitting the PROTAX model

PROTAX converts sequence similarities into probabilities of taxonomic classification in a hierarchical manner, starting from the root node of the taxonomy and proceeding towards the species nodes. Each node divides its probability into its child nodes by means of a multinomial regression model. The predictors used in the multinomial regression can be chosen in many ways. While the results of Somervuo et al. (2016) suggest that a combination of similarity-based and phylogenetic-based predictors yields the best performance both for simulated and real data, in this study we used solely similarity-based predictors.

The regression model for each taxonomic node containing seven predictors *x*_1_,…,*x*_7_. The baseline case where all the seven predictors are zero corresponds to a child node that represents a missing branch of the taxonomy. Predictor *x*_1_ is an indicator variable for a known child node that contains no reference sequences, whereas predictor *x*_2_ is an indicator variable for a known child node that contains at least one reference sequence. Predictors *x*_3_ and *x*_4_ are, respectively, the mean and the maximum value of pairwise sequence similarities between the query sequence and the reference sequences. To allow PROTAX to account in the predictions for the availability of the number of reference sequences (with which e.g. maximal similarity is expected to increase just by chance), we included as predictors also the log-transformed number of reference sequences representing the child node (*x*_5_), and the interactions between log-transformed number of reference sequences and mean (*x*_6_) and maximal (*x*_7_) similarities.

We calculated pairwise sequence similarities using LAST (Kielbasa et al. 2011) with the following deviations from the default parameters. We set the LAST argument -T 1 to make the similarity score represent the entire overlap alignment length between two sequences, excluding only the possible overhangs. We set the gap open penalty to (-a 1). In order to get meaningful values to the mean sequence similarity predictor of the PROTAX model, we set the maximum number of initial matches per query position (-m) values between 1000 and 3000 instead of the default value 10. We replaced pairwise sequence similarities that were missing from LAST output by zeros, and converted sequence similarities to the range [0,1] by dividing the alignment score by the alignment length.

We generated training data to parameterize the PROTAX model as described in Somervuo et al. (2016), i.e. by modifying both the taxonomic tree itself as well as its coverage by the reference sequences to mimic the different kinds of outcomes: (i) known species with reference sequences, (ii) known species without reference sequences, and (iii) unknown species or unknown higher taxonomic branches. For each case study, we generated in total 1000 training data points, out of which 100 represented the category (iii), with an even distribution over the taxonomic levels. The remaining 900 sequences representing categories (i) and (ii) were generated by randomly selecting one of the species present in the database, and generating training data directly for that species, or if not possible, for another species that was taxonomically as close to the selected species as possible. For example, if the selected species had no reference sequences, we selected the closest species that had at least one such sequence, selected one sequence to represent the query sequence, and removed all the other sequences to mimic a species with no reference sequences.

In our baseline analyses, we assumed a priori that all species that are part of the Linnaean taxonomy are equally likely to be present in the empirical sample. If there is prior information about which species are more likely to be found from an empirical sample than others, such information can be incorporated into PROTAX by a weighting scheme, which can be considered as an informative prior in the context of Bayesian analyses. To illustrate the influence of the prior, we conducted an alternative version of the fungal analyses, where we gave more prior weight for those species that are known to occur in Finland, as our environmental samples originate from there. From the list of Finnish 6645 fungal species, we could map 4718 names to the 130 795 species taxonomy. In the weighted analysis, we assumed a priori that each sequence present in our environmental sample represents one of the species known to occur in Finland with probability 90%, and thus dividing the remaining probability of 10% among the remaining species.

We derived maximum *a posteriori* (MAP) parameter estimates for the PROTAX models using the Bayesian approach presented in Somervuo et al. (2016), except that in the present study we parameterized the models separately for each taxonomic level. The model parameters for each level include the seven regression coefficients corresponding to each of the predictors, as well as the probability by which the reference sequence is mislabeled (Somervuo et al. 2016).

### Evaluating the performance of PROTAX

We used the parameterized PROTAX models to perform taxonomic placements of both reference sequences as well as environmental sequences. In the first set of analyses, we performed taxonomic placements for 1000 validation sequences, which were chosen from the reference sequence database in the same way as the training sequences described above. While PROTAX yields for each of these the full probability distribution over possible outcomes, we selected here only the outcome with the highest probability. We considered a taxonomic placement as “plausible” if the classification probability was at least 50%, and as “reliable” if the classification probability was at least 90%. To examine the overall confidence of classifications, we computed the proportions of plausible and reliable classifications at each taxonomic level. To assess if the probabilities of taxonomic placement were unbiased, we ordered the classification probabilities from lowest to highest, and computed a cumulative sum of both these probabilities as well as the indicator variables describing whether the outcome predicted with highest probability was a correct one. We then plotted these two cumulative sums against each other. If the classification probabilities are unbiased, such a plot should follow the identity line.

In the second set of analyses, we performed taxonomic placements for the environmental sequence data. As the mammalian, fungal and bacterial case studied involved a large number of sequences generated by high-throughput methods, we first clustered these sequences. The data submitted to PROTAX involved 1514 (mammal), 4163 (fungi), and 6855 (bacteria) OTUs, and 7939 individual insect sequences.

To visualize community composition within each environmental data set, we used Krona (Ondov et al. 2011) to plot for each case study a pie chart that shows the expected number of sequences representing each taxonomic unit. To compute the expected abundances, we did not account only for the highest probabilities, but we summed over the entire distribution of predicted probabilities (ignoring values lower than 0.01 for computational reasons). To visualize the quality of the classifications, we colored the charts to show six categories. The first three categories consisted of well-identified taxonomic units for which the proportion of sequences for which the classification was reliable was (1) in the range 50%-100%, (2) in the range 0%-50%, or (3) 0%. The remaining three categories consisted of non-identified taxonomic units for which the proportion of sequences for which the classification was reliable was (4) in the range 50%-100%, (5) in the range 0%-50%, or (6) 0%. Above, well-identified taxonomic unit refers to a single taxonomic unit for which reference sequences were available, whereas non-identified taxonomic units refers to the union of taxonomic units without reference sequences and unknown branches of the taxonomy.

## Results

As expected based on our earlier results (Somervuo et al. 2016) and the fact that PROTAX is a statistical model fitted to training data, PROTAX yielded essentially unbiased probabilities of taxonomic placement for all the cases considered. This is evidenced by the fact that all lines in Fig. 2 generally follow the identity lines, the small deviations being attributable either to sampling error due to finite sizes of the validation data sets, or to issues related to model misspecification, the latter of which we return to in the Discussion. The probabilities shown in Fig. 2 are level-specific, thus asking e.g. how well genera can be separated within a known family, or how well species can be separated within a known genus. For high taxonomic levels, these probabilities are lowest for fungi, which is consistent with the fact that for fungi there is the greatest amount of overlap in sequence similarities among consecutive taxonomic levels (Fig. 1). For example, if within-species similarities are sometimes lower than among-species similarities, accurate taxonomic placement to the species-level is not always possible.

**Figure 2.**
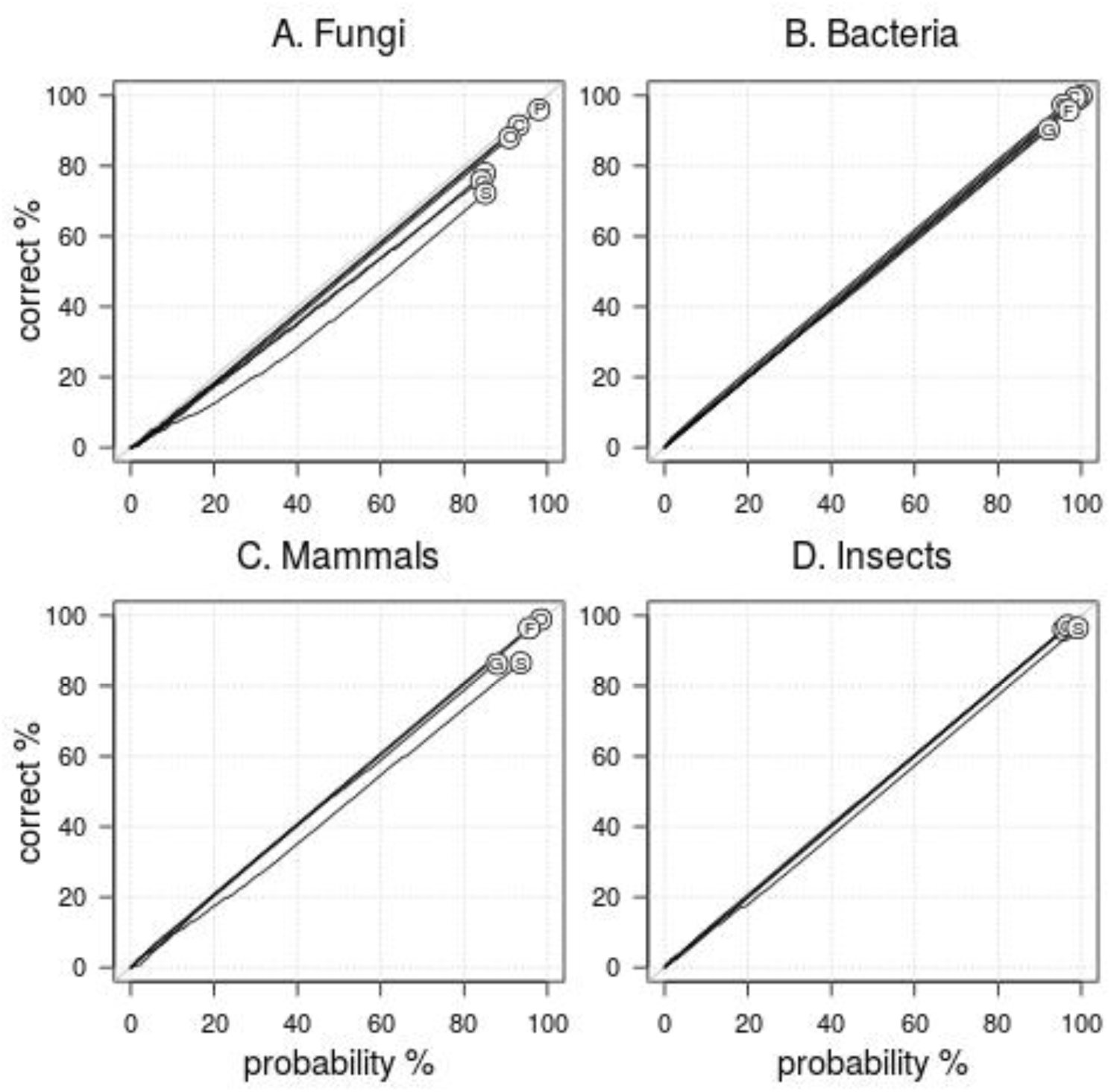
An assessment of bias and accuracy of the PROTAX algorithm for classifying well-identified sequence data to different taxonomic levels. We used PROTAX to classify well-identified reference sequences, with the focal sequence removed from the reference database to avoid circularity. The classification probabilities shown here are level-specific conditional probabilities, thus measuring e.g. the accuracy of species-level classifications conditional on knowing the true genus. While PROTAX yields a vector of identification probabilities for all possible outcomes, we considered here only the outcome with the largest identification probability, which we compared to the true identity of the species. For each taxonomic rank (indexed as S=species, G=genus, F=family, O=order, C=class, D=domain), panels show the cumulative number of correct identifications on the y-axis versus the cumulative sum of the identification probabilities on the x-axis (both normalized by the number of sequences). A curve matching with the identity line (y=x) indicates unbiased identification probabilities, both for small and large probabilities, as the identifications have been sorted in the order of increasing largest identification probability. The position of the dot gives the mean identification probability among the samples.

When performing a taxonomical placement of environmental samples, PROTAX works in a hierarchical manner starting from the root of the tree, and proceeding level by level towards the tips of the tree that represent typically species. The probabilities of taxonomic placement for a given level (illustrated in Figs. 3 and 4) are thus obtained by multiplying the level-specific conditional probabilities (illustrated in Fig. 2) for all levels lower than or equal to the focal level. Figure 3 shows the proportions of the reference sequences (black lines) and environmental sequences (gray lines) that were possible to identify reliably (dashed lines) or plausibly (continuous lines). Let us first make two obvious remarks. First, as the threshold for plausible identification (>50% probability of taxonomic placement) is lower than that of reliable identification (>90% probability of taxonomic placement), the proportion of plausible identifications is always higher than that of reliable identifications. Second, as the lower level taxonomic placements are conditional on the higher level ones, the fraction of reliable (and plausible) identifications decreases monotonously with taxonomic level.

**Figure 3.**
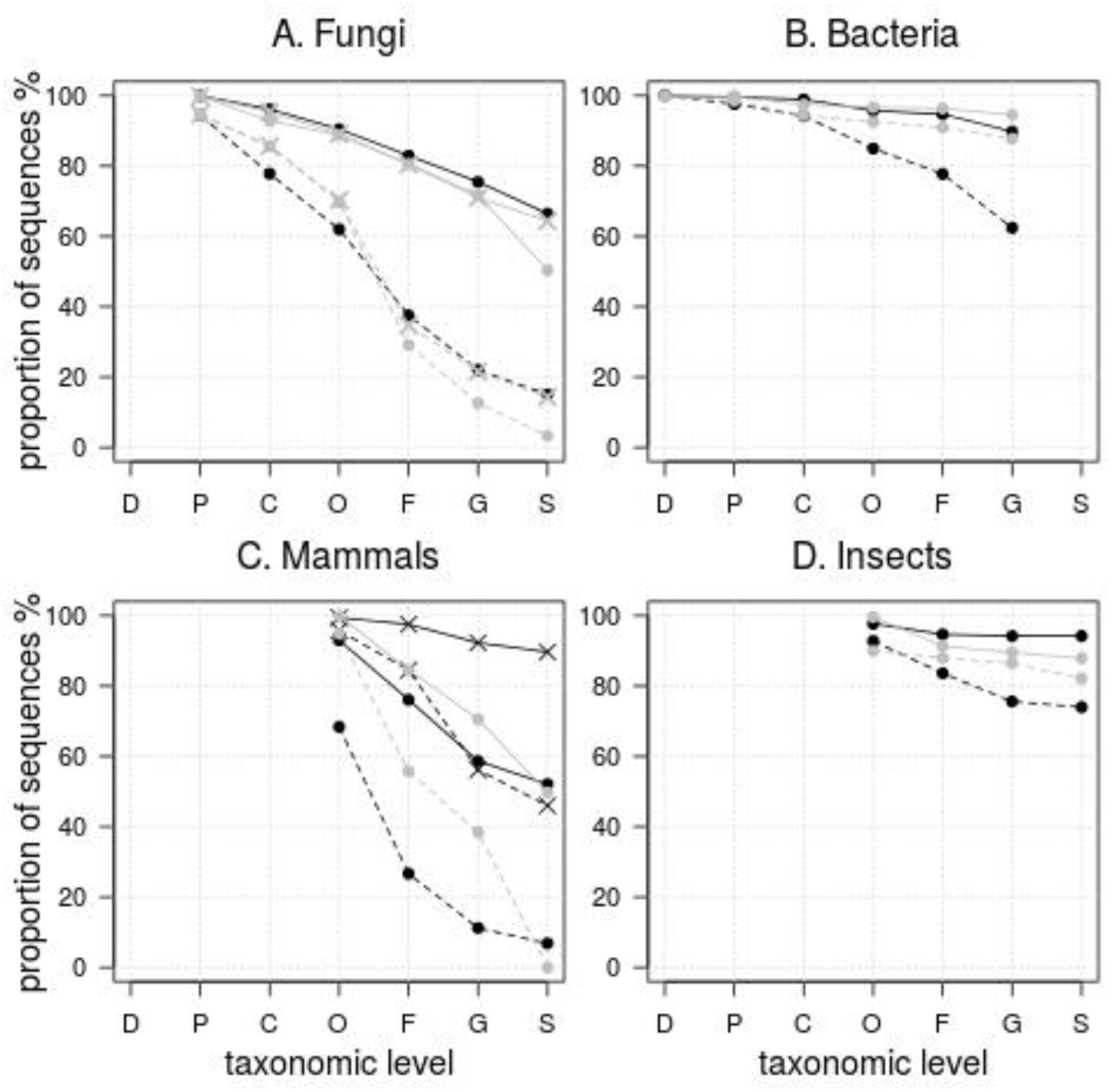
Confidence of taxonomic placement at different taxonomical levels. The value on the y-axis is the proportion of plausible (solid line) and reliable (dashed line) taxonomic placements. Results for validation data sampled from reference sequence database are shown in black and results for environmental query data are shown in gray. For fungi, gray crosses denote results from environmental data where species probabilities were weighted according to prior knowledge on which species exist in Finland. For mammals, black crosses denote results using full-length mt 16S rRNA sequences as validation data. Taxonomic labels at x-axis from left to right: D=domain, P=phylum, C=class, O=order, F=family, G=genus, S=species.

**Figure 4.**
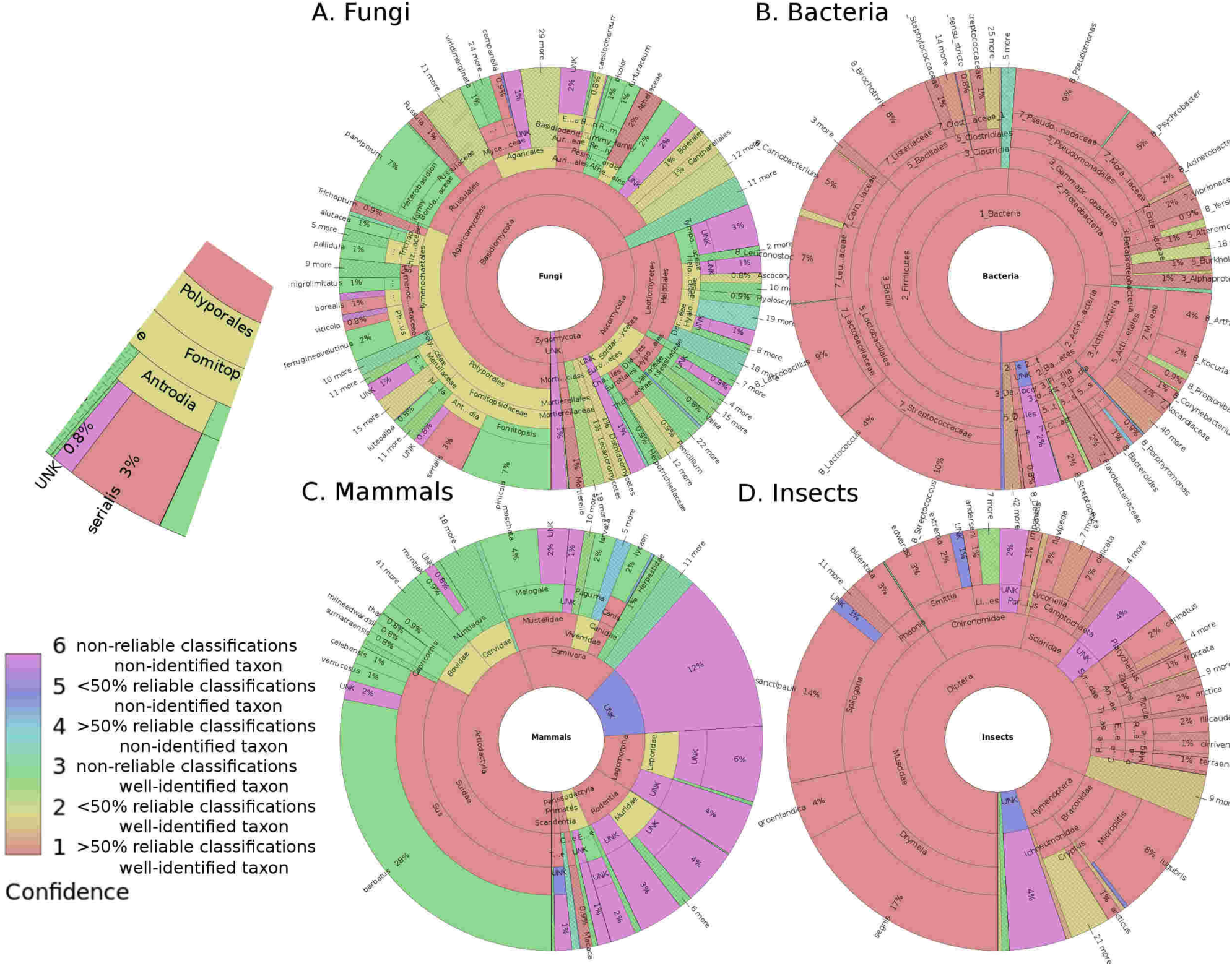
Taxonomy pie charts of PROTAX output showing the composition of the environmental data sets. The width of each sector is proportional to the expected number of sequences that was placed to that taxonomic units. The colors code both the reliability of the identifications, and whether the identifications relate to taxonomic units that are part of the taxonomy or to unknown units (see color label). The enlarged insert illustrates species-level resolution for the fungal data. The charts are snapshots from interactive web pages (provided in the Supporting Information) generated by Krona software from the PROTAX output.

Beyond the above made trivial remarks, Fig. 3 shows a number of interesting results. As the first result, that we derive from the taxonomic placement of the validation sequences, reliable species-level identification (dashed black lines in Fig. 3) was most successful for insects (74% of the sequences), followed by mammals (46%) and fungi (15%). These numbers do not reflect only the resolution of the barcoding sequences (Fig. 1), but also the fact that the insect taxonomy and reference sequence databases were restricted to species occurring in Greenland, whereas the mammalian and fungal databases were global and thus were larger and more heterogeneous (Table 1). For mammals, full-length mt 16S sequences (black crosses in Fig. 3C) can be expectedly classified with much higher confidence than fragmented sequences (black dots in Fig. 3C), the latter corresponding to the nature of the environmental data. In case of bacteria, reliable genus-level identification was possible for the majority (62%) of the cases.

As the second result, Fig. 3 shows that taxonomic placement of environmental sequences is often less reliable than that of reference sequences (mammals and fungi), but sometimes environmental sequences can be identified essentially equally reliably (insects) or even more reliably (bacteria) than reference sequences. The main reason why taxonomic placement of environmental sequences for mammals was much more difficult than that of reference sequences is simply that in our case study the environmental sequences were very short fragments. If fragmenting the reference sequences equally much (into 100 bp segments), their taxonomic placement became essentially equally unreliable than that of reference sequences (lines with black dots in Fig. 3C). In case of fungi (Fig. 3A), the reason for the difference between the taxonomic placement of the reference and environmental sequences was not only a similar (though less pronounced) difference in sequence length and quality as for mammals, but also the fact that the environmental sequences are likely to represent many unknown units that are lacking from the taxonomy. If bringing the prior information that, instead of any globally known fungi, the species within the environmental sample are likely to represent species that are known to occur in Finland, the proportion of reliable identifications increases dramatically from 3% to 14% (Fig. 3C). The reason why for the insect data (Fig. 3D) the taxonomic placements are essentially equally reliable for the reference and environmental sequences is that for this case study both kinds of sequences were acquired by identical methods, i.e. Sanger sequencing of DNA sampled from individual specimens. Thus, the only differences between the two were whether the specimens were identified morphologically or not, and whether the specimens represent a random sample of the community (environmental sequences) or whether they were targeted to represent the entire community (reference sequence data). The most curious case is that of bacteria, where reliable genus level taxonomic placements were more frequent for environmental sequences than for reference sequences (Fig. 3B). The likely reason here is that in this case the environmental sequences originated from the food production pipeline, the bacterial communities of which represent one of the most well studied groups, and thus are better covered in the reference sequence database than bacteria in general.

Let us then turn into the main question that motivates DNA (meta)barcoding studies: what are the species behind the environmental samples? The answer to this question is given in Fig. 4, where the pie charts show the proportions of sequences that belong to known and unknown taxonomical unit at each hierarchical level. In this figure, the areas of the sectors show the expected number of sequences that belong to each taxonomic unit, whereas the colors illustrate the proportions of reliable identifications, and they thus echo the information shown by the grey dashed lines in Fig. 3. While our main interest here is not on the detailed results relating to the four case studies, let us note that the overall patterns in Fig. 4 are consistent with expectations. Concerning fungi, the majority of the species Agaricomycetes, and the reliably identified species (e.g. *Antrodia serialis;* see the insert in Fig. 4) typically represent well known wood decomposers. Concerning mammals, both Artiodactyla, Chiroptera, Rodentia and Carnivora were detected, as well as some primates. While there are very few reliable or even plausible species-level taxonomical placements, among possibly identified species are e.g. the endangered mammals *Muntiacus vuquangensis* (Giant Muntjac; 43% identification probability) and *Rusa unicolor* (sambar; 27% identification probability). Concerning bacteria, a large proportion of the sequences were assigned as Lactobacillales, specifically to Streptococcaceae, Lactobacillaceae, and Leuconostocaceae (Figure 4). Further, the high proportion of *Brochotrhrix* observed by Hultman (2015) was supported by the PROTAX results. Concerning insects, the majority of the species belonged to Diptera and the minority to Hymenoptera. Among the total of 104 distinct species that were reliably identified, the most common one was *Drymeia segnis*, which has been observed to be common in the study area also based on morphological identifications (Rasmussen et al. 2013).

In Supporting Information, we provide the same information as shown in Fig. 4 as interactive HTML files, which allow the pie charts to be displayed using a standard web browser without any additional plugins. This allows one to examine the taxonomic placements and their reliabilities in much greater detail by e.g. using search tools and zooming to taxonomic clades of specific interest.

## Discussion

In this work, we have evaluated the potential of DNA barcoding for obtaining reliable taxonomic placements at different taxonomic levels, and in particular illustrated how the PROTAX method can be used as a general tool for quantifying uncertainty in such taxonomic placements. PROTAX accounts for many kinds of uncertainties, including the possibilities of unknown taxonomic branches, incomplete coverage of reference sequence databases, and mislabelling of reference sequences. This makes its quantification of taxonomic placement uncertainty robust, as illustrated by Fig. 2 and the simulations by Somervuo et al. (2016). However, it is important to understand that the classification accuracy does not necessarily increase when taking all uncertainties into account; it can rather be the opposite. To put it bluntly, it may be more tempting e.g. to claim that the study detected the endangered mammal Giant Muntjac from a leech blood meal, rather than to specify that this was the case with 43% probability, as the latter statement makes it explicit that the species behind the sequence may actually have been some other one. However, making uncertainty explicit is necessary for scientific reliability.

There are many choices to be done when applying DNA (meta)barcoding to an empirical case study. As illustrated by our results, these choices can have a major influence on the reliability of the resulting taxonomic placements. The first set of choices relates to the taxonomy and reference databases used, which choices are in practice mostly guided by on what databases are available rather than what might be optimal to use. Importantly, as PROTAX accounts for missing branches in the taxonomy, the incompleteness of the taxonomy database should not lead to spurious false positives, rather to decreased probabilities of taxonomic placement. This is because in the training phase PROTAX generates situations in which some branches of the taxonomy are missing, making it learn which kinds of values of the predictors (e.g. low values of sequence similarity) are indicative of missing branches. Similarly, mislabeled reference sequences or inconsistencies between the taxonomy and the reference databases are expected to decrease the probabilities of taxonomic placement, but not to bias them. As one example, we used the RDP database for bacteria. Since the reference taxonomy was constructed based on the reference sequences, 100% of the taxa in the validation data were covered (Table 1). Somewhat surprisingly, the bacterial reference database appeared to represent also the vast majority of the environmental sequences, with only very few missing branch identified (Fig. 4). This however does not mean that the used taxonomy would cover all the bacteria in the world, and novel phyla have indeed been discovered in several recent metagenomic studies (e.g. Brown et al. 2015). The other commonly used bacterial and archaeal databases are SILVA (Quast et al. 2013) and Greengenes (DeSantis et al. 2007). Compared to RDP, these two databases contain more representatives of the Candidate divisions that have been recently found in various soil environments (Brown et al. 2015; Hug et al. 2016). Therefore, depending on the environment under analysis, the use of different reference databases should be considered.

The second set of choices to be made relates to the DNA barcode applied, as well as the sequencing technology. As has been long pointed out, an optimal barcoding gene should involve much variation among species but only little within a species (Meyer and Paulay 2005). Further, the environmental sequences should obviously have as long read length and as high quality as possible. For example, if in the mammalian case study full length mt 16S rRNA sequences had been available instead of the very short 100 bp fragments used here, the proportion of reliable taxonomic placement would have been likely to increase from the present 0% to ca. 46%, where the latter was the proportion of reference sequences that we could classify reliably. But even if one would have full length sequences and complete taxonomic and reference sequence information, some uncertainty will inevitably remain. For example, in the insect study the mosquito species *Aedes impiger* and *Aedes nigripes* could not be disentangled since their COI sequences are identical, and thus PROTAX assigned for some of the specimens a probability close to 50% for both of these species. To resolve such cases, a deeper genomic approach (Bourke et al. 2013) than the single gene DNA barcoding approach should be used.

The third set of choices relates to the way in which the training data in PROTAX are generated, technically the prior assumed for the empirical data. This is probably the most critical and at the same time most difficult choice to be done by the user, as making a justified choice requires biological knowledge and intuition. For example, one may assume either that each sequence in the environmental sample represents any of the species present in the taxonomy with equal probability (as we have done here), or utilize a hierarchical prior that assumes that each branch under a given node is equally likely (as we did in Somervuo et al 2016). One may further give additional weight for species that are known to occur in the geographic region where the samples originated, as we did for the fungal case study. If such information is available, the prior can also be adjusted e.g. based on the expected abundances of the species, or on the match between the substrates sampled and the habitat requirements of the species. In addition to the known species, the prior involves an assumption about the frequency of missing branches at different parts of the taxonomic tree. As it may be difficult to make informative choices about all of the above mentioned aspects, we recommend the user the test the sensitivity of the results against different choices of the prior, as should be done with Bayesian analyses in general.

Finally, the fourth set of choices relates to the predictors used for the multinomial regression underlying the PROTAX model. In this paper, we have used simply similarity-based predictors, even if our previous work suggests that similarity-based predictors and phylogeny-based predictors involve complementary information and thus their combination optimizes performance (Somervuo et al. 2016). The reason behind the choice made for the present work was mainly computational, as some of our databases were extensive, making LAST-based similarity the most practical choice. For fungi, the use of phylogeny-based predictors is challenging also for the reason that the construction of multiple sequence alignments is difficult with the ITS region only. Phylogeny-based methods are easier/more suitable with conserved barcodes such as CO1 and mt16S which allow sequences to be globally aligned even at high taxonomic levels. In more refined studies focusing on any specific case study, the set of predictors should be optimized to maximize the reliability of taxonomic placements. While there is no objective way to select the best prior, the choice of the predictors can be optimized more or less objectively by examining which predictors maximize unbiased probabilities of taxonomic placement for independent validation sequences. The reason why for some choices of the predictors the classification probabilities can be biased (as was to a limited extent a case for some of our case studies, Fig. 2) is that while the PROTAX model is parameterized by training data, the model may be structurally misspecified. For example, we have assumed that the model parameters are constant across the taxonomic tree. Thus, when classifying an environmental sequence e.g. to the species level under a known genus, the parameters (and thus the influences of the predictors, such as sequence similarity) are assumed to be independent of the genus. This assumption is not likely to hold for large and heterogeneous taxonomic groups, such as all mammals or all fungi. An indication of this in our results was that, at the species level, the parameter estimates obtained for mislabeling probability were much inflated, being ca. 80% for mammals and ca. 60% for fungi. This does not suggest that there is such a vast amount of mislabeling, but that PROTAX used the mislabeling parameter to correct for model misspecification. Thus, an important challenge for future work is to further develop the statistical model underlining PROTAX, either by building a hierarchical structure that allows for heterogeneity in the parameterization, or by finding predictors that are able to correct for such heterogeneity.

To conclude, molecular species identification by DNA barcoding and metabarcoding is an exciting and rapidly evolving research field, which has major potential to change our understanding of the structure and functioning of ecological communities. To make the use of these methods practical and reliable, a key challenge is the completion and pruning of taxonomic and reference sequence databases, as well as making these two sources of information compatible. Similarly important is the application and further development of statistical methods that allow one to make the most out of such data by providing accurate taxonomic placements and reliable assessments of the uncertainties inherent in such placements. Such methods are critical for providing a firm basis for deriving species- and community-level inferences from DNA (meta)barcoding data, especially for environmental DNA that by definition do not have physical specimens that could be verified independently. Incorrect assignments can result in accumulated interpretation error, which can result in wasted resources and social conflict in multiple social arenas, from conservation to food safety. It is important to get the name right – or to be aware that it may be wrong.

## Acknowledgements

We thank Mikko Tiusanen and Tomas Roslin for providing the empirical sequence data for the insect case study, and Paul Kirk and Kessy Abarenkov for providing the taxonomy and reference databases for the fungal case study, and Otto Miettinen for providing the list of Finnish fungal species. OO and PS were supported by funding from the Academy of Finland (Grants no. 1273253 and 250444 to OO) and the Research Council of Norway (CoE grant no. 223257). DWY and YQJ were supported by WWF-Vietnam (9S084701), the National Natural Science Foundation of China (31400470, 41661144002), the Ministry of Science and Technology of China (2012FY110800), the University of East Anglia and its GRACE computing cluster, and the State Key Laboratory of Genetic Resources and Evolution at the Kunming Institute of Zoology (GREKF13-13, GREKF14-13, and GREKF16-09). CX was supported by the MEME Erasmus Mundus Programme in Evolutionary Biology, and the Marco Polo Exchange Fund of the University of Groningen.

